# DeXtrusion: Automatic recognition of epithelial cell extrusion through machine learning *in vivo*

**DOI:** 10.1101/2023.02.16.528845

**Authors:** Alexis Villars, Gaëlle Letort, Léo Valon, Romain Levayer

## Abstract

Epithelial cell death is highly prevalent during development and in adult tissues. It plays an essential role in the regulation of tissue size, shape, and turnover. Cell elimination relies on the concerted remodelling of cell junctions, so-called cell extrusion, which allows the seamless expulsion of dying cells. The dissection of the regulatory mechanism giving rise to a certain number and pattern of cell death was so far limited by our capacity to generate high-throughput quantitative data on cell death/extrusion number and distribution in various perturbed backgrounds. Indeed, quantitative studies of cell death rely so far on manual detection of cell extrusion events or through tedious systematic error-free segmentation and cell tracking. Recently, deep learning was used to automatically detect cell death and cell division in cell culture mostly using transmission light microscopy. However, so far, no method was developed for fluorescent images and confocal microscopy, which constitute most datasets in embryonic epithelia. Here, we devised DeXtrusion, a pipeline for automatic detection of cell extrusion/cell death events in larges movies of epithelia marked with cell contour and based on recurrent neural networks. The pipeline, initially trained on large movies of the *Drosophila* pupal notum marked with fluorescent E-cadherin, is easily trainable, provides fast and accurate extrusion/cell death predictions in a large range of imaging conditions, and can also detect other cellular events such as cell division or cell differentiation. It also performs well on other epithelial tissues with markers of cell junctions with reasonable retraining.

## Introduction

Epithelial tissues can be dramatically remodelled during embryogenesis or in adult organs undergoing fast turnover. This is often associated with high rates of cell elimination, which requires the fine control of the absolute number of dying cells, as well as their distribution in time and space. Cell extrusion is a sequence of remodelling steps leading to apical constriction and cell elimination from the epithelial layer without impairing the sealing properties of the tissue (Rosenblatt et al., 2001). This process is highly coordinated between the extruding cell and its neighbours: as the cell extrudes its neighbours are brought close to each other to maintain epithelial sealing and stability (Villars and Levayer, 2022).

The tight spatiotemporal and quantitative control of epithelial cell apoptosis plays an essential role during tissue morphogenesis (Ambrosini et al., 2017). For instance, a spatial bias in the distribution of cell death can locally modulate growth and final tissue shape (Matamoro-Vidal et al., 2022), apoptosis and cell extrusions can generate local traction forces to fuse tissues (Toyama et al., 2008), promote locally tissue bending (Monier et al., 2015; Roellig et al., 2022) or can be permissive for global tissue remodelling through the modulation of tissue viscosity (Ranft et al., 2010; Suzanne et al., 2010). Moreover, the tight regulation of the precise spatiotemporal distribution of cell extrusion/cell death is also essential to maintain the cohesion of the tissue, especially in conditions with high rates of cell elimination (Valon et al., 2021). These examples all rely on the precise regulation of the number and spatiotemporal localisation of dying cells. Yet, despite the fast progress in our understanding of the molecular regulators of programmed cell death and extrusion, we still fail to predict when, where and how many cells will die in tissue. This most likely relies on the multiple feedback that can modulate death rate/cell extrusion at various spatial and temporal scales (Villars and Levayer, 2022), which includes the activation of the prosurvival signal ERK in the neighbouring cells (Bock et al., 2021; Gagliardi et al., 2021; Valon et al., 2021), long-range coordination for cell extrusion (Aikin et al., 2020; Takeuchi et al., 2020) as well as positive feedbacks on apoptosis (Perez-Garijo et al., 2013). Thus, obtaining a comprehensive understanding of epithelial cell death regulation entails the dissection of multi-layered regulations integrating feedback on several spatial and temporal scales as well as several steps of decisions. Such a challenging question requires a highly quantitative dataset on the total number of cell death as well as their precise spatiotemporal distribution and high throughput methods to compare these values in various perturbed backgrounds.

The recent advances in long-term live imaging provide a wealth of data regarding tissue dynamics, especially for epithelia in 2D. However, retrieving cellular events quantitatively in a high throughput manner remains extremely challenging. For instance, there are more than 1000 extrusions, distributed all over the epithelium in the *Drosophila* pupal notum in less than 12h (Guirao et al., 2015; Valon et al., 2021). So far, quantitative analyses of cell death/extrusion were performed using laborious manual detection of these events (Moreno et al., 2019; Valon et al., 2021; Villars et al., 2022). This highly time-consuming task remains one of the main bottlenecks for comparing a high number of conditions with precise quantitative readouts. Alternatively, automatic epithelial cell death detection was performed through systematic segmentation and tracking of all the cells (Etournay et al., 2016; Guirao et al., 2015). However, this method typically entails extensive manual corrections even when using machine learning-enhanced segmentation (Aigouy et al., 2020), and still represents quite an important load of work for large fields of view and long timescales, hindering large-scale analysis.

Altogether, these challenges and needs call for an automatic tool that would allow accurate spatiotemporal detection of cellular events without relying on systematic segmentation and tracking of cells. The recent progress of computer vision opened the possibility to automatise the detection of objects and patterns in biological images and image series (Hallou et al., 2021). In particular, deep learning approaches were used successfully to recognise cellular events such as cell division and cell death in yeast (Aspert et al., 2022) or in mammalian cell culture (Kabir et al., 2022; La Greca et al., 2021; Mahecic et al., 2022; Phan et al., 2019; Shkolyar et al., 2015). However, this was mostly applied to transmission light microscopy and these pipelines are not applicable to large samples and embryos, where imaging mostly relies on fluorescent and confocal microscopy. As such, there is currently to our knowledge no implemented solution for the automatic detection of cell death and cell extrusion events from epithelia *in vivo*.

To answer these challenges, we devised here a supervised machine-learning pipeline called DeXtrusion. The detection of extrusion is performed by screening sliding windows spanning the entire movie and detecting cellular events on each window. The core of our pipeline is based on a recurrent neural network which classifies each image sequence. These local classifications are then post-processed together at the movie level to convert them to probability maps of the presence of a cellular event or to single point event detections. We devised and applied this method on the *Drosophila* pupal notum, a single-layer epithelium, using cell-contour-labelled epithelia with tagged E-cadherin. The method is flexible enough to provide accurate predictions with movies of different temporal resolutions, pixel sizes, imaging set-ups as well as different E-cadherin labelling. Moreover, DeXtrusion generalises well out-of-the-box to other epithelia (e.g.: pupal abdomen, pupal wing), which can be further enhanced by retraining using small training sets. By resolving the bottleneck of extrusion/cell death detection, DeXtrusion will open the way for a more systematic and quantitative characterisation of cell death distribution in large datasets which will be essential to understand its multi-layered regulation. DeXtrusion is distributed as a python module, available open source on gitlab (https://gitlab.pasteur.fr/gletort/dextrusion) along with our trained neural networks and scripts (jupyter notebooks and Fiji macros) to facilitate its usage, our annotated datasets used for training of the pipeline are available on Zenodo (Villars et al., 2023) (https://doi.org/10.5281/zenodo.7586394).

## Results

### DeXtrusion: a neural network to recognise extrusion in cropped image sequences

We first aimed at detecting cell extrusion in the *Drosophila* pupal notum: a single-layer epithelium on the back of the developing *Drosophila* pupal thorax with a high rate of cell extrusion/apoptosis that follows stereotypical patterns (**Fig.1A**) (Guirao et al., 2015; Levayer et al., 2016; Marinari et al., 2012; Valon et al., 2021; Villars et al., 2022). We used a dataset generated in the laboratory covering a large number of extrusion events (6700 in the training set and 2320 in the test set see **Supplementary tables 1 and 2**). This dataset was made of large-scale movies of the pupal notum obtained from two different imaging set-ups (see **Methods**), with different frame rates, signal-to-noise ratios, and different fluorescent proteins coupled to the adherens junction protein E-cadherin. It contains both Wild Type (WT), mutants and drug-perturbed conditions to obtain a model robust in a large range of conditions (see **Supplementary tables 1 and 2**). Cell extrusions were manually annotated while control positions were drawn randomly, excluding locations containing an extrusion. To handle data with different frame rates and spatial resolutions, we fixed a reference spatiotemporal scale (0.275 microns/pixel and 5 min/frame) on which the neural network was trained.

**Figure 1:**
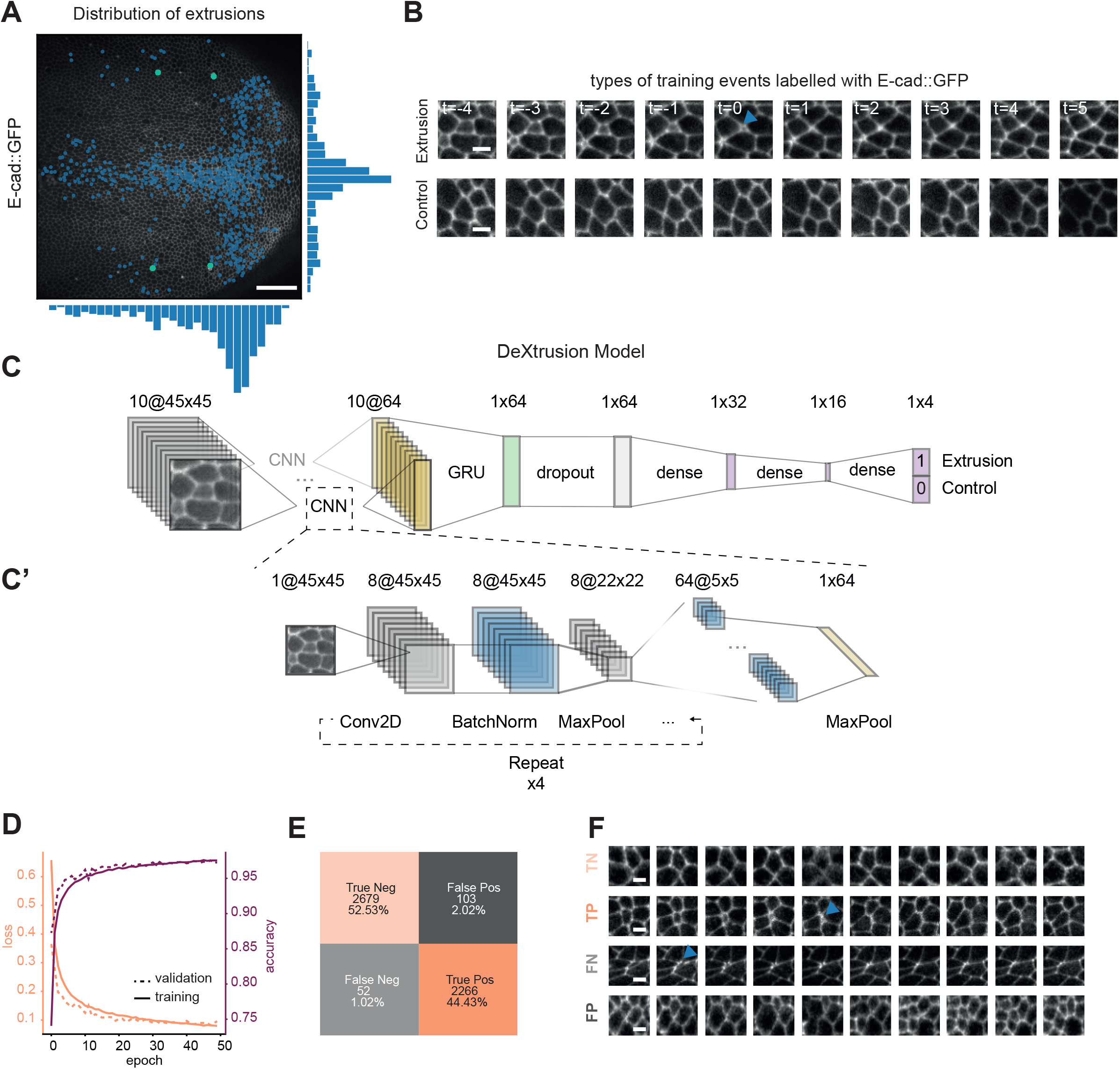
DeXNet, a neural network to recognise cell extrusions in image sequences in epithelia. **A:** Snapshot of a developing *Drosophila* pupal notum marked with E-cad::GFP (Knockin) with overlaying extrusions manually detected over 16h of development. Histograms represent the distribution of cell extrusion along the antero-posterior axis (bottom histogram) and along left-right axis (right histogram). Scale bar=50 μm. **B:** Representative image sequence centered on an extrusion event (top row, blue arrow points at the end of extrusion), or representing a control sequence with no event (bottom row). The time is given in frame (here, one frame =5 minutes). Scale bar=5 μm. **C-C’:** Schematic of the model architecture called DeXNet. **C.** Main model. The image sequence is passed first to a CNN detailed in **C’** which encodes the image in a 10×64 matrix which is itself fed into a Gated Recurrent Unit (GRU) to take into account temporal information. This is then passed to a dropout normalisation layer before going through a sequence of densely connected layer (dense). Finally, it predicts with a probability whether the input sequence is an extrusion or not. **C’:** Detailed architecture of the encoding CNN. Each image in the sequence goes into a sequence of convolution (Con2D), Batch normalisation and max pooling (MaxPool, all this is repeated 4 times) before going through a final step of max pooling encoding the image in a final vector (1×64). **D:** loss (left y-axis) and accuracy (right y-axis) curves for training data (bold lines) and validation data (dotted lines). **E:** Confusion matrix showing the accuracy of the DeXtrusion model. Orange coloured boxes show the number of correctly predicted events (light orange are True Negatives, darker orange show True Positives). Grey events are the number of Wrongly predicted events (light grey are False Negative and darker grey show False Positives). **F:** Representative image sequences showing example events for each of the categories in **E**(TN: True Negative — Event correctly classified as controls, TP: True Positives — Extrusions events correctly classified as extrusions, FN: False Negative — Extrusions wrongly classified as control events, FP: False Positives — Control events wrongly classified as extrusions). Scale bar=5 μm.

To detect with spatiotemporal accuracy all cell extrusions, we generated a pipeline called DeXtrusion, that screened through the entire movie using overlapping sliding windows. Each cropped image sequence (a series of 10 images of 45×45 pixels in the reference scale) was run through a neural network (DeXNet) to estimate the presence or absence of an extrusion event (**Fig.1B**). For this, we built and trained a neural network to classify each image sequence (**Fig.1C-C’**). We first chose an architecture based on Convolutional Neural Network (CNN), efficient for image-based classification (LeCun et al., 2015). However, since CNN on single images performed poorly at predicting extrusion, we decided to include temporal information by using a recurrent neural network architecture. We chose a Gated Recurrent Unit (GRU) architecture which computes and propagates temporal information while preserving a parsimonious network architecture (Cho et al., 2014). Each image of the temporal sequence is first encoded using the CNN (**Fig.1C’**) and reduced in a vector of relevant features, which is then combined with the other image feature vectors into the GRU (**Fig.1C**). The output of the time distributed GRU layer then goes through a sequence of densely connected layers that perform the final classification (**Fig.1C purple**). Eventually, the network provides a probability of containing an extrusion for each cropped image sequence. This probability was finally thresholded to a binary output (extrusion or control event). We trained the DeXNet on image sequences extracted from our dataset, split in a training and test set (25 versus 7 movies, see **Methods**). We took cropped image sequences in the reference scale centred spatially and temporally around the manually detected extrusions (time 0 being defined by the termination of apical constriction), while control sequences were made of similar cropped image sequences that do not contain any event (see **Methods**). The temporal size of the sequences was chosen so that it contains enough information to recognise extrusion while reducing the chance to capture two events (see **Methods**, **Fig. S1A-C**). The x-y size of the cropped input images, as well as other hyperparameters of the model (number of epochs -the number of times the dataset is processed by the network for training-, of features, data augmentation…), were tuned by testing several sets of hyperparameters (**Fig. S1, S2 and S3**). Taken together, the selected hyperparameters allowed the model to converge properly under 50 epochs and took around 55 min of training with 1 GPU (**Fig. 1D**). Compared to our manual annotation, the classification of the neural network on our test dataset resulted in a precision of 0.97, with 2% False Positive and 1% False Negative (**Fig.1E-F**). As our neural network performed well at spotting cell extrusions in cropped image sequences, we then integrated it into the DeXtrusion pipeline to detect all the cell extrusions in a large-scale movie.

### From cropped images classification to whole-movie extrusion detection

So far, our trained neural network DeXNet provides a fast and efficient classification of each image sequence. To detect extrusion events across the tissue and at all temporal time points, the complete movie was split into 45×45 pixel^2^ cropped images with 50% overlap. The time axis was divided into sequences of 10 frames every two time points after rescaling the time to the reference scale (**Fig. 2A**). These parameters are a good compromise between the spatiotemporal precision of the results and the speed of computation and can be tuned by users. For each sliding window, DeXNet provides a single probability of extrusion. This value is then allocated in the original movie in a 22×22 pixels and 5-time points regions around the centre of the cropped sequence. The final local probability output is then obtained by averaging for each pixel the probability of the different overlapping windows (in time and space) generating an extrusion probability map on the whole movie (**Fig. 2A,B, Movie 1**). This output can be directly used to screen visually the detection of extrusion events and assess putative tissue patterns (**Fig. 2B**).

**Figure 2:**
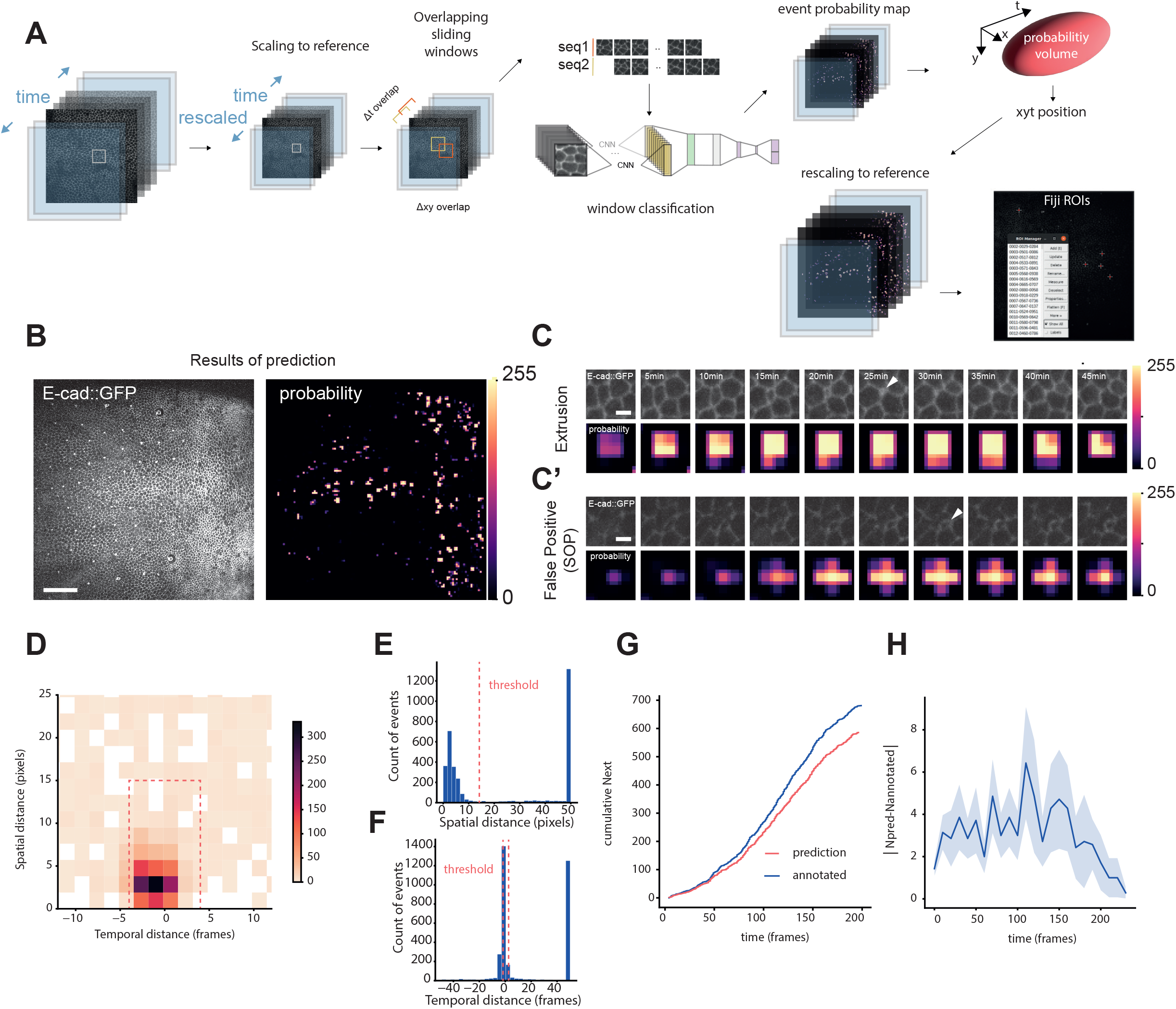
DeXtrusion pipeline to detect extrusions on full movies. **A:** Schematic representation of the DeXtrusion pipeline. The input movie is first rescaled to the reference scale. Then, cropped image sequences are extracted, spanning the entire movie with an overlap both spatially and temporally. Each image sequence is then processed through our neural network DeXNet. The resulting probability of the presence of an extrusion is added around the central position of the image sequence to build a probability map on the entire movie, before rescaling it to the original movie size. Single point events can also be generated by taking the centroids of high probability volumes in the probability map and exported as a Fiji ROI file. **B:** Projection of a resulting probability map. Snapshot of the input movie, an epithelium labelled with E-cadherin-GFP (left) with the corresponding probability map (right). Probabilities are drawn as a color map, with values converted to 0 (black) - 255 (white) scale for visualisation. Scale bar=50 μm **C:** Example of correctly detected extrusion and of False Positive detection. Image sequences of E-cadherin labelled epithelium original movie cropped around a correctly (top) or wrongly (bottom) detected extrusion event (top line). Corresponding image sequences of the resulting probability maps (below). Probabilities are color coded from 0 (black) to 255 (white). White arrows indicate detected event. Scale bar=5 μm. **D:**2D histogram of the spatio-temporal distances between manually annotated extrusions and DeXtrusion results on the 7 test movies. The colour code represents the number of extrusions detected within a given temporal distance (x-axis, in time frames, 1 frame=5 minutes) and a given spatial distance (y-axis, in pixels, 1 pixels=0.275μm). The red dotted rectangle represents the events that are considered as matching with a manually annotated event (below the spatial and temporal threshold, see **Methods**). **E:** Histogram of the distribution of the spatial distance of DeXtrusion results to the closest manually annotated extrusion. The dotted line represents the spatial threshold used to associate the detected extrusion with a manually annotated extrusion. The peak of detections at 50 pixels corresponds to extrusions that do not match temporally (more than 4 time frames distance). **F:** Histogram of the distribution of the temporal distance of DeXtrusion results to the closest manually annotated extrusion. The dotted line represents the temporal threshold used to associate the detected extrusion with a manually annotated extrusion. The peak of detections at 50 frames correspond to extrusions which do not match spatially (further than 15 pixels). **H:** Cumulative number of extrusions during developmental time for DeXtrusion results (red) and manually annotated one (blue). Extrusions were detected on one test movie of 942*942 pixels and 200 frames. **I:** Average absolute difference of the number of extrusions over time between DeXtrusion detections and manually annotated extrusions. From all test set movies (binned every 10 min from the start of the movie).

To count the number and precisely localise cell extrusion events, this probability map must be converted to point-like event detection. Our overlapping method detects single events within several consecutive time windows and on a given x-y surface, thus generating a volume (x-y-t) of pixels with high probability (**Fig. 2A,C, Movie 1**). We first thresholded the probability map to obtain a 3D mask of positive detections. To filter out false positives which are usually detected on smaller areas and fewer time windows (**Fig. 2C**), we also thresholded the size of the positive volume to fit the minimal volume associated with extrusion events (which last between 20 and 30 minutes (Villars et al., 2022)). The probability threshold and volume threshold were optimised to maximise the precision and recall (**Fig. S2, Methods**). We also used a watershed separation to separate close events. Eventually, the exact location of the extrusion is defined by the centroid of the high-probability volume. Detected extrusions are then exported as a list of point ROI (Region Of Interest) compatible with Fiji ROI manager (Schindelin et al., 2012) (**Fig. 2A**).

To assess the performance of DeXtrusion, we compared the resulting ROIs with the manually annotated ROIs of our test dataset. We first measured systematically the x-y Euclidean distance and time distance between manually annotated and automatically detected extrusions (**Fig. 2D-F**). This showed that 70.6% of the detections are below a 15-pixels distance and 4 time frames shift. We then classified detections as correct for spatial distance below 15 pixels (~ 1 cell radius) and below 4 time-frames (+/- 20 minutes) distance between the DeXtrusion and manually located ROIs. Doing so, we obtained a recall of 0.87 (proportion of events detected) and a precision of 0.46 (proportion of detected events that are indeed extrusion). Finally, we measured systematically the total number of detected extrusions compared to manually annotated events at every developmental time to check whether the accuracy of our detection is sensitive to the developmental stage. We observed a fairly constant error (~3 errors every 10 frames/50 minutes) suggesting that our methodology is robust at all developmental stages imaged in the notum (**Fig. 2G-H**). Thus, our methodology can retrieve the vast majority of the extrusion events at any stage of development.

### Optimisation of the model for the detection of extrusion

So far, we have designed and optimised DeXtrusion to detect exhaustively all the extrusion events, which led to the high recall of 0.87 (87% of events detected). However, we have not optimised the methodology to filter out false positive detections. Indeed, many cellular events which share some phenotypic similarities with extrusion are very frequently miss labelled as extrusion (**Fig. 3 A-D**, **Movie 1**). This includes for instance the formation of Sensory Organ Precursors (SOPs) which through asymmetric cell division form cells with very small apical area (Gho et al., 1999) which are very often detected as extrusion (**Fig. 3B**). Similarly, the shrinkage of cell apical area concomitant with cytokinesis and furrow formation are frequently misclassified as extrusion (**Fig. 3C**). Thus, to enhance the precision of DeXtrusion (the proportion of correct extrusion detection), we decided to add these other cellular events in DeXNet to discriminate them from extrusion. We trained the network to detect cell extrusions, cell divisions and SOPs or the absence of events using manually annotated events in our dataset (6700 extrusions, 3021 divisions and 3054 SOPs, **Fig. 3F**). DeXNet was able to classify the cropped image sequences for these 4 events with high accuracy on our test set (**Fig. 3G**, **Movie1**). By including this new DeXNet in our pipeline, we significantly increased the precision of DeXtrusion (from 0.26 to 0.41, **Fig. 3E,E”**), while this did not impact significantly the recall (**Fig. 3E’**). This is also reflected by the F1 score (**Fig. 3E**), a measurement of precision and exhaustivity of detections (see **Methods**from 0.40 to 0.54). To further enhance our precision, we then manually screened all remaining false positive detections and included image sequences representative of the patterns of these false positives in the training inputs as control sequences. This reinforcement increased the model’s precision to 0.52 for similar recall (0.92 compared to 0.90). Finally, we exploit the inherent stochasticity of the training process (which always lead to different network weight and biases) to confront the predictions from two independently trained networks (Segebarth et al., 2020). Averaging these two independent classifications had the most significant effect and increased the precision to 0.72 (recall of 0.86 and f1-score of 0.78). This barely affected the recall (**Fig.3E-E”**, 2nets), but it increased the calculation time by a factor of two. Of note, the accuracy was much lower for the test movie of the colcemid injected pupae (**Fig. 3E-E”, deep blue, movie ID18**), which phenotype was poorly represented in the training dataset and corresponded to an extreme condition where the mode of extrusion and the architecture of the tissue are quite different (Villars et al., 2022). After all these optimisation steps, the final model (2nets) can still detect all cellular events on a movie covering the full pupal notum (1200 by 1200 pixel^2, 200 time points) in less than 40 min on a regular PC with 1 GPU. Taken together, these optimisations steps led to a drastic increase in precision while maintaining similar recall on test movies. The last remaining false positive (~20%) can be easily filtered out manually from the final ROIs list. We estimated that the full procedure (DeXtrusion computation time+manual correction) takes between 1 or 2 hours for a movie covering the full pupal notum over 200 times (representing roughly 1000 extrusions). This corresponds to a drastic time saving compared to the exhaustive manual detection of extrusions (~10 hours for a trained user (Valon et al., 2021) versus approximatly one hour of manual correction with DeXtrusion).

**Figure 3:**
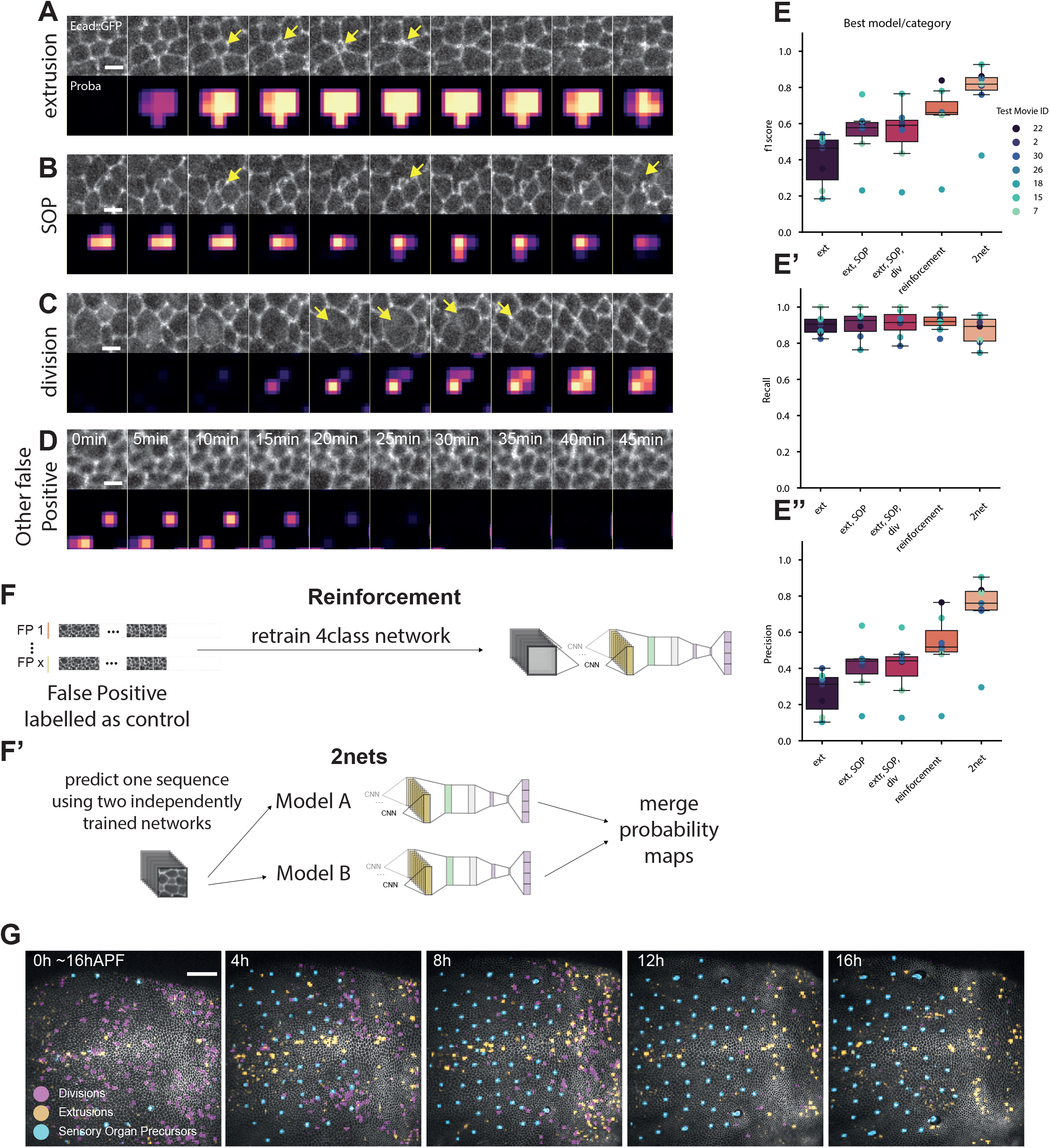
Optimisation of the model for the detection of extrusion. **A-D:** Image sequence events predicted as extrusion correctly (**A**) or not (**B-D**) by the initial DeXnet trained on 2 classes only (extrusion or not). Top rows of each panel represent the image sequence, bottom rows represent the probability maps obtained by DeXtrusion. Scale bar= 5 μm. **A:** Image sequence representing an extrusion and its associated probability map. Yellow arrows point at the closure of the extruding cell. **B:** Image sequence representing a forming Sensory Organ Precursor (SOP) wrongly predicted as an extrusion and its associated probability map. Yellow arrows point at the small cell of the SOP which constricts and leads to the misclassification. **C:** Image sequence representing a dividing cell wrongly predicted as an extrusion and its associated probability map. Yellow arrows point at the furrow formation during cytokinesis. **D:** Image sequence representing a control event wrongly predicted as an extrusion and its associated probability map. **E-E”:** Changes in the model to optimise its prediction scores on extrusion. **E:** f1-score. **E’:** Recall, **E”:** Precision, with the initial two class model (ext), the inclusion of SOPs (ext, SOP), the inclusion of SOPs and cell divisions (ext, SOP, div), including the 3 cellular events and reinforcement (see below), and using two independent networks (2net). The results are shown for the best trained network for each class model (see **Fig. S2 G-I** for the averaged). **F-F’:** Schematics of the steps added to the model to increase its f1 -score on predicting extrusions. **F:** Reinforcement consists in taking control events misclassified as extrusion and to add them to the training set with a control label. This forces the model to learn that the previously misclassified events are in fact controls. **F’:** Models are stochastically trained which results in models with slightly different weight and biases. Using the average of the two best trained models increases the f1 -score and the robustness of the prediction. **G:** Image sequence showing the results of DeXtrusion predictions after optimisation, of extrusions (orange), cell divisions (pink) and differentiation (SOPs, blue) on full-scale movie (pupal notum, local projection of E-cad::GFP) and overtime. Time is shown in hours After Pupal Formation (hAPF). Scale bar= 50 μm.

### Generalisation of DeXtrusion to mutant context and other epithelia

The performance of a neural network is usually highly limited to the training data range (Mockl et al., 2020), which may also apply to DeXtrusion. To test the generalisation of our model, we therefore, challenged our pipeline by testing its performance on different data from closely similar biological and imaging conditions in very different contexts. We first trained two “full” networks, using all our annotated data (training+test dataset). To adapt DeXtrusion to other tissues (**Fig. 4A**) where the cell size and duration of cellular events are not of the same scale, the reference scale at which movies are resized is calculated so that cell diameter is around 25 pixels and extrusion taking place over 4-5 frames. We first tested DeXtrusion on *Drosophila* pupal notum depleted for EGFR (UAS-EGFR-dsRNA driven by pnr-Gal4), a condition that modifies tissue shape and the spatiotemporal distribution of extrusion while not affecting so much the extrusion process per se (Moreno et al., 2019; Valon et al., 2021). We obtained an overall good detection level in this context (**Fig. 4A,B,B’**,**E Movie 2**). Since annotated data do not exist for these movies, we randomly sampled the prediction results to manually check the events (see **Methods**). Using this method, we estimated that 87% of extrusion detections were correct (precision=0.87, **Fig. 4E**). We further challenged DeXtrusion using the *Drosophila* pupal wing, an epithelial with similar cell shape and size to the pupal notum (Aigouy et al., 2010; Etournay et al., 2015; Farhadifar et al., 2007) (**Fig. 4A,C,C’**, **Movie 3**). Using a previously published movie of E-cad::GFP pupal wing (Etournay et al., 2015), we obtained a precision of 0.786 (excluding ROIs outside the wing, see **Methods**, **Fig. 4E**) and an estimated recall of 0.85 (estimated on 59 extrusions manually annotated). These scores could be significantly improved by retraining DeXNets on a small proportion of manually annotated extrusions and divisions from the pupal wing (75 events), reaching a precision of 0.91 and an estimated recall of 0.83, **Fig. 4E**, **Movie 3** and **Methods**).

**Figure 4:**
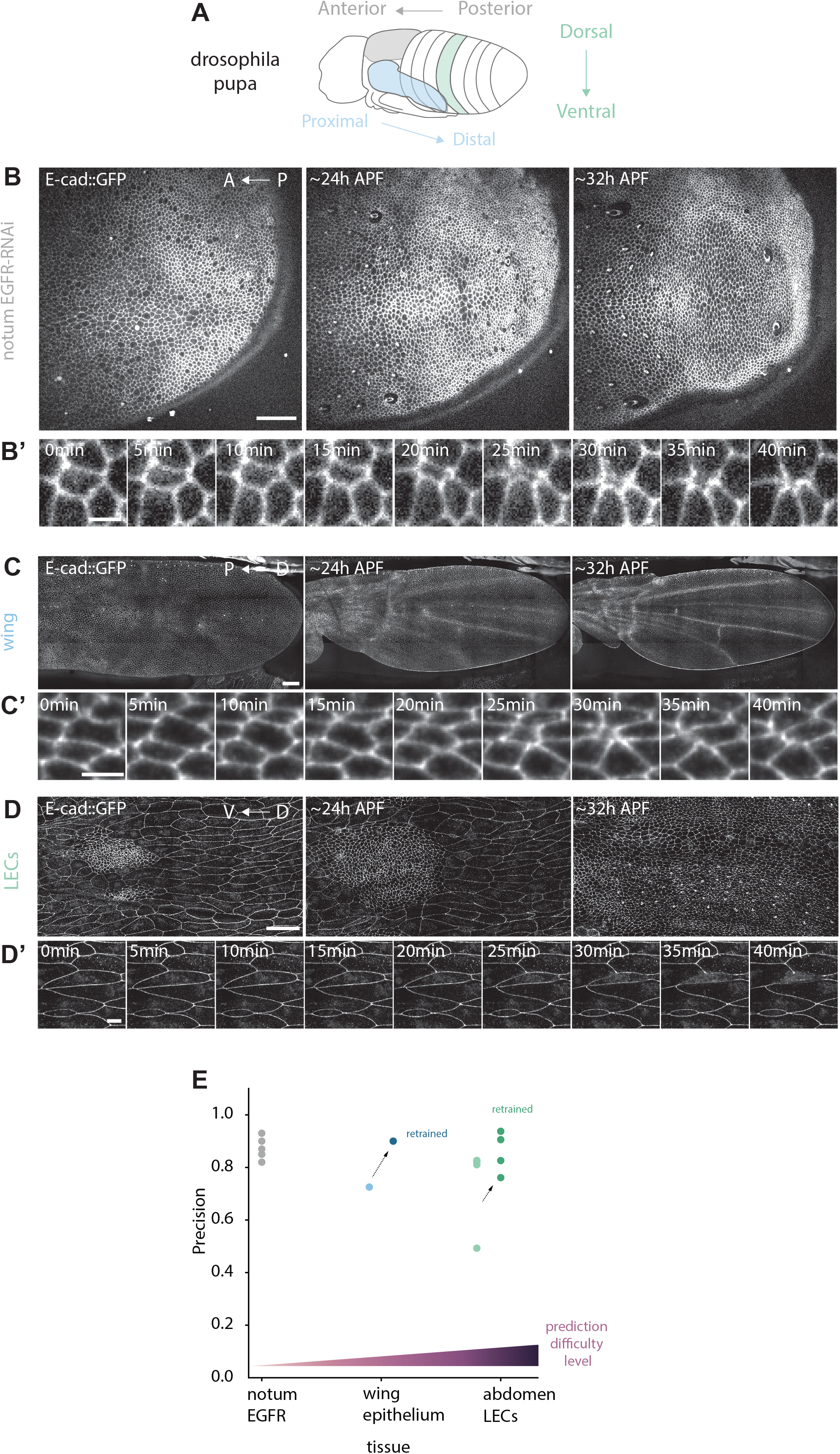
Generalisation of DeXtrusion to mutant context and other epithelia. **A:** Schematic of a *Drosophila* pupa highlighting the different epithelia on which the model was tested. The model was trained on the notum (green) and tested on the pupal wing (blue) and finally on the abdominal Larval Epithelial Cells (LECs, magenta). **B-D’:** Example image sequences of the different tissues used to test the generalisation of DeXtrusion and the corresponding extrusion phenotypes. **B:** Pupal notum epithelium expressing a RNAi against EGFR. Scale bar=50 μm. **B’:** Example image sequence showing an extrusion cropped from **B**. Scale bar=5 μm **C:** Pupal wing epithelium from (Etournay et al., 2015). Scale bar=50 μm. **C’:** Example image sequence showing an extrusion from **C.** Scale bar=5 μm. **D:** Pupal abdomen epithelium extracted from (Davis et al., 2022), results were computed only for LECs (bigger cells) excluding the histoblasts (smaller cells). Scale bar=50 μm. **D’:** Example image sequence showing an extruding LEC from **D**. Scale bar=10 μm. **E:** Manually computed precision for the prediction on the different tissues with increasing differences compared to the tissues for which DeXtrusion was trained on. Green shows the precision of DeXtrusion for notum expressing a RNAi against EGFR (example shown in **B-B’**). Light blue is the precision of DeXtrusion for the pupal wing epithelium (example shown in **C-C’**). Dark blue is the precision of DeXtrusion on the same tissue after retraining the model on a subset of extrusions. Light purple is the precision of DeXtrusion on LECs (example shown in **D-D’**). Purple is the prediction on the same tissue after retraining. Darker purple is the prediction on the same tissue after retraining and reinforcement.

We further challenged DeXtrusion by testing its performance on a squamous epithelial, the larval epithelial cells of the pupal abdomen, where cells have very different shapes from the pupal notum and where extrusions occur through slightly different mechanisms (Hoshika et al., 2020; Michel and Dahmann, 2020; Teng et al., 2017; Villars et al., 2022) (**Fig. 4 A,D,D’**). This test was conducted on 4 movies from (Davis et al., 2022; Tapon and Salbreux, 2022) focusing exclusively on the larval epidermal cells. After proper rescaling, we obtained a precision of 0.737 but obtained a low recall of 0.38 (using available segmentation and annotation (Davis et al., 2022; Tapon and Salbreux, 2022), despite adjusting the threshold distances (spatial and temporal) to the scale of the movie (see **Methods**). This may reflect the difference in cell morphology (long and curved junctions) and dynamics of extrusion (progressive loss of E-cad and rounding, (Teng et al., 2017), **Fig.4 D’**). To adapt DeXtrusion to these cells, we retrained our DeXNet models using the annotated extrusions of one of the four movies. The precision increased to 0.857 and the recall nearly doubled to 0.69 (**Fig. 4E**, **Movie 4**). Increasing the retraining data to 2 movies continued to improve the performance but less drastically (precision of 0.86 and a recall of 0.74).

Altogether, this demonstrates that DeXtrusion can robustly detect extrusion events on various tissues and conditions. For situations where cell shape and the profile of extrusions are very different, few additional trainings are sufficient to reach back good precision and recall, illustrating the adaptability of DeXtrusion.

## Discussion

In this study, we have designed a fast tool that allows the automatic detection of several cellular events and estimated its accuracy. While the pipeline was primarily optimised for the pupal notum with labelled E-cadherin, the same principle can easily be applied to any other epithelial tissue provided there is a minimal dataset to retrain the model. While our precision (~0.8) still requires a manual correction phase, this leads to a considerable gain of time compared to the manual annotation of large movies (from 10 hours to 1 hour) which could lead to rapid quantification of a large number of movies. This considerable gain of time opens new opportunities for systematic quantification of the spatiotemporal distribution of cell death in a large numberof movies. DeXtrusion offers the possibility for large screening of drugs/mutations on the spatiotemporal distribution of cell death that could lead to the identification of new biological factors modulating cell apoptosis and new spatiotemporal feedbacks. While we extended DeXtrusion to detect cell division and SOPs with the aim of improving our precision, these additional features can also be handy to analyse the spatiotemporal interplay between these three cellular events, for instance regarding the coupling between cell death and cell division (Fan and Bergmann, 2008; Kawaue et al., 2021; Mesa et al., 2018). Note however that we did not estimate the accuracy of our pipeline to detect these other cellular events as it was not the initial scope of our study. We challenged DeXtrusion to detect extrusions in other tissue and demonstrated how easily it could be adjusted to detect new extrusion patterns or cellular junction staining. We believe it could easily be trained to learn to recognize new cellular events as well. In particular, a significant proportion of false positives were related to local cell rearrangements in the tissue, related to junction remodelling and T1 transitions (Guirao and Bellaiche, 2017). Adding this new feature could not only increase the precision of our pipeline but also offer an additional feature for large-scale analysis of tissue dynamics.

We distributed DeXtrusion as an open-source pipeline with a graphical interface to propose an efficient and user-friendly tool to the community. Our main aim was to reduce drastically the necessary manual input to allow large-scale analysis of cell extrusions in the *Drosophila* pupal notum. The final tool allowed to go further by offering the possibility to screen also for cell divisions and SOPs and moreover can be easily adapted to add other cellular events or to other kinds of cellular labelling.

## Acknowledgements

We would like to acknowledge the Image Analysis Hub of the Institut Pasteur for discussions and advice for this work. We would also like to thank Raphaël Etournay for sharing the pupal wing movie and Nic Tapon for providing the original and segmentation of the pupal abdomen movies. We acknowledge the help of the HPC Core Facility of the Institut Pasteur for this work. AV was supported by a PhD grant from the doctoral school “Complexité du Vivant” Sorbonne Université and from an extension grant of La Ligue contre le Cancer, work in RL lab is supported by the Institut Pasteur (G5 starting package), the ERC starting grant CoSpaDD (Competition for Space in Development and Disease, grant number 758457), the ANR CoECECa, and the CNRS (UMR 3738).

## Authors contributions

AV and GL contributed equally to that work. AV gathered the dataset, designed the model and conducted initial testing. GL revised the model and the code and extended it on the prediction of full movies. Both AV and GL contributed equally to the figure and results. LV contributed to the dataset with his data. AV, GL and RL wrote the manuscript and designed the project.

## Methodology

### Generation of the training dataset

To create an annotated dataset, we used movies that had been previously manually annotated for extrusions in the laboratory for other studies (Moreno et al., 2019; Valon et al., 2021; Villars et al., 2022). Pupae were dissected and imaged on a confocal spinning disc microscope (Gataca systems) with a 40X oil objective (Nikon plan fluor, N.A. 1.30) or 100X oil objective (Nikon plan fluor A N.A. 1.30) or a LSM880 equipped with a fast Airyscan using an oil 40X objective (N.A. 1.3), Z-stacks (1 μm/slice). All the movies were built using a local z-projection plugin which follows tissue curvature (Herbert et al., 2021). We pooled together 32 movies, all from the *Drosophila* pupal notum, but with different acquisition setups, genetic backgrounds and junctions staining. The full list of movies and their characteristics is given in **Supplementary tables 1 and 2**. We split the annotated data into two datasets, one for training (**Supplementary table 1**) and a smaller one for testing (**Supplementary table 2)**. For this, we selected randomly one movie of each of our different conditions for the test set, and all remaining movies of the same condition were used for training. We obtained two datasets: the training set of 25 movies, with 6692 annotated extrusion events from 8 different conditions, and a test set of 7 movies, with 2320 annotated extrusions from 7 different conditions. These data with the manual annotation are freely available on a repository (https://doi.org/10.5281/zenodo.7586394) (Villars et al., 2023).

For each network training, the training dataset was split in 75-25% between training and validation data. Note however that these two subsets were not fully independent as they are composed of windows extracted from the same movies (training movies).

### Training image sequence generation

To generate training image sequences from the ROI files and not keep all the movies in the running memory, we implemented a movie generator (MovieGeneratorFromRoi.py). It randomly selects the ROI from the input file and saved a cropped image sequence centred around that point, with a small spatial and temporal random shift (so that the event is not perfectly centred in the window to limit a bias toward the centre of the image). For control (no event) windows, positions are randomly drawn in the movie and kept if there is no ROI in it.

### Data augmentation

#### Movie augmentation

The movies acquired at higher spatial and temporal resolution are much smaller compared to the majority of movies once rescaled in the reference scale. As such, this kind of data was under-represented in the training data, with only a few events by movies. To reduce this bias, we doubled these movies by downsampling temporally each original movie two times and introducing a shift in the frames extracted between the two repetitions. These movies are indicated with “_aug” at the end of the names in the available dataset.

#### Image sequences augmentation

We also performed data augmentation on the training cropped image sequences. To augment the generalisation of the training, the augmentation was done on each training window with the addition of small temporal and spatial shifts, gaussian noise, white/black squared, and illumination on all windows or one time-frame.

### Data imbalance

The events that we classified (nothing, extrusion, division, SOP) do not have the same frequency in all movies. To reduce the possible imbalance between the representation of these events in the training data, we reduced the number of windows of each over-represented event used in the training to have a number of training data similar than the less represented events. This option can be turned off in the pipeline with the boolean parameter “balance”.

### Data rescaling

To homogenise the training dataset and have events of similar duration/size, we rescaled all our dataset to the same temporal and spatial resolution of 5 min/frame and 0.275 μm/pixel. Rescaling of the movies and ROIs was done with a Fiji macro. DeXNet networks were trained with input windows of this reference scale. A division event was visible on 2 to 3 time-frames, cell extrusion on 4 to 5 frames, and a cell had a typical diameter of 25 pixels. Therefore, the DeXtrusion pipeline must rescale all the movies to this reference scale before applying the classification. Since characteristic event duration and cell sizes can vary between tissues/organisms, we used these “cellular” features for rescaling rather than absolute time or distances.

### Data to test generalisation

We tested the robustness of DeXtrusion to different unseen datasets. The first dataset was composed of 5 movies of *Drosophila* pupal notum depleted for EGFR (UAS-EGFR-dsRNA) (Valon et al., 2021). We do not have manual annotations on these movies, so they were not used in the training data. However, 3 movies in the original dataset were acquired with the same imaging and genetic conditions. This dataset was thus considered very similar to the training data.

The second dataset was a large movie of the *Drosophila* pupal wing from (Etournay et al., 2015) (3879×1947 pixels, 200 time-frames). This tissue was not represented in the training data, but the organisation of the epithelium and cell shape is very similar to the pupal notum. This dataset was considered similar to training data. We manually annotated a few extrusions (59) spanning the movie to evaluate the recall of DeXtrusion in this sample. We extracted a small part of the movie (495×444 pixels and 200 frames) and annotated this cropped movie to use as retraining data with 27 extrusions, 29 divisions and 19 additional controls for reinforcement. Note that the field of view of the movie contains the whole wing but also external tissue on the top and bottom parts. We focused the quantification only on the ROIs that were fully inside the wing.

The third dataset was composed of four movies of larval epithelial cells (LECs) of the *Drosophila* pupal abdominal (Davis et al., 2022). The annotation of cell extrusions in the larval epidermal cells was obtained using the segmentation mask provided in the original dataset (Tapon and Salbreux, 2022) through tissue miner (Etournay et al., 2016). We focused our test only on larval cells and excluded histoblasts (the nest of small cells) since there is hardly any extrusion in this population in an attempt to focus on cells different from the pupal notum. We estimated the typical cell diameter of the larval cells to 80 pixels and the extrusion duration to 10 time-frames and used these values to rescale the movies for the pipeline. To estimate the Recall, we compared DeXtrusion outputs with the generated extrusion annotations on the original movie. However, since cell size and extrusion durations were much higher in these movies, we adjusted the thresholds of spatial and temporal distances to count matching detection (50 pixels xy distance, ~half a cell, and 8 time-frames). Note that the scores were always calculated on the 4 movies, even when some were used in the retraining data which could induce a slight positive bias. However, the 2 movies used for retraining were the ones on which DeXtrusion results were best even before retraining, and the most drastic improvements came from the 2 other movies.

### DeXtrusion source code

DeXtrusion is available open-source on gitlab at https://gitlab.pasteur.fr/gletort/dextrusion under the BSD-3 license. The trained neural networks, Jupyter notebooks and Fiji macros to use DeXtrusion are available and described in this repository. DeXtrusion main code is deployed as a python module that can be installed through the pip installer package: https://pypi.org/project/dextrusion/. Instructions to install and use DeXtrusion are given in our gitlab repository.

To ease its usage, we proposed Jupyter notebooks that are dedicated to tasks such as network training, retraining or DeXtrusion detection on new movies. To visualize the results as probability maps or ROIs, Fiji macros are also available. To handle input/output between our python code and Fiji, we used two specific python modules in our pipeline: “roifile” (Gohlke, 2022a) and “tifffile” (Gohlke, 2022b).

### DeXNet architecture and training

To categorize sliding windows by taking into account temporal information, we based the architecture of our neural networks on the Gated Recurrent Unit (GRU) architecture (Cho et al., 2014). We tested different variations of the architecture and hyperparameters (number of layers, number of features by layer, number of epochs, size of the sliding windows…). The final architecture is represented in **Fig. 1C** and the full detailed architecture can be found in the source code in the Network.py file. The parameters used for training one network are summarised in the configuration file associated with each DeXNet. For training the neural network to categorise the window as containing an event or not, we used the categorical cross-entropy loss.

In our Gitlab repository, we proposed 2 DeXNets trained for controls and extrusions classifications (notum_Ext, used for **Fig. 1** and **Fig. 2**), controls, extrusions and SOPs classifications (notum_ExtSOP, **Fig. 3**), controls, extrusions, SOPs and cell divisions classifications (notum_EXTSOPDiv, **Fig. 3**), and trained on all events and all data (train and test) pooled together (notum_all, used for **Fig. 4**).

### Evaluation of DeXtrusion results

#### DeXNet evaluation

The performance of DeXNet networks was measured by the accuracy of the results during training, and with the confusion matrix of the classifications on the test dataset after training (**Fig.1E**).

#### Pipeline evaluation

##### Comparison with manual annotations

To estimate the quality of DeXtrusion detections, we computed the precision (TP/(TP+FP)) and recall (TP/(TP+FN)) of the resulting ROIs compared to manual annotated ROIs (TP: True Positive, FP: False positive, FN: False Negative). ROIs were considered to be the same (between results and manual annotations) if they were within a spatial distance of 15 pixels and a temporal distance of 4 frames (in the reference scale). The Jupyter notebook dextrusion_CompareRois.ipynb allows us to calculate these scores and to choose the threshold distances to consider ROIs as the same. To consider both precision and recall at once, we also measured the F1-score (TP/(TP+(FP+FN)/2)) to evaluate the performance of our pipeline.

##### Measure of False Positive without manual annotations

For movies on which we do not have manual annotations, we cannot measure the recall as this would necessitate full annotations of all the events. We estimated the percentage of False Positive detections by selecting randomly a high number of resulting ROIs and examining each ROI manually to decide if the hit was correct or not. The Fiji macro deXtrusion_scoreROIs_Random.ijm allows to do it and gives the resulting percentage.

##### Model Optimisation

Once the model architecture was fixed, we optimised most of the model parameters to achieve the best possible results on the prediction of extrusion. For this, we started with the optimisation of the exploration time window (**Fig. S1**) as it is the first input of the model. The training gave the best results when the exploration window was temporally centered on extrusion (5 frames before, 5 frames after) (**Fig. S1**). We then used this parameter to assess the effect of the window’s xy size. The best results were obtained for a half size of 21px. While this parameter doesn’t yield the best precision (**Fig. S1D**), it gives the best recall out of all parameter values explored (**Fig. S1E**). Recall was the metric we tried to optimise the most to avoid missing any extrusion. Moreover, the prediction time increased linearly with the window size (**Fig. S1F**). As a result, once pondered by time (precision*recall/prediction time), a size of 21px appears as the best value (**Fig. S1G**).

We then explored the impact of the different model thresholds on the model results (**Fig. S2**). We first fixed a volume threshold of 800px to assess the impact of the probability threshold (**Fig. S2A-C**). While the best f1-score is obtained for a probability threshold of 200, it comes at the expanse of a lower Recall (**Fig. S2A & C**). Thus, we set up to use a value of 180 for that parameter (second best but higher Recall, **Fig. S2A**) to explore the impact of the volume threshold and following similar reasoning picked a value of 800 for that parameter.

Finally, we tried to optimise the hyperparameters (parameters of the model for training) (**Fig. S3**). First, we explore hyperparameters during the training of a model predicting only 2 classes (extrusions vs nothing). After parameters exploration we selected an augmentation of 3 (**Fig. S3A**), a number of epochs of 40 (**Fig. S3B**), a number of CNN filters of 8 (**Fig. S3C**) and a number of reinforcements of 5 (**Fig. S3D**). We then optimised the model by adding new predicting classes (SOPs and division, **Fig. 2**) and thus applied the same approach to the model prediction with 4 classes (extrusion, SOPs, divisions, nothing). Changing the number of epochs had a very limited impact and we, therefore, kept a number of 40 epochs for the final model and an augmentation of 2.

All selected parameters are summarized in **Supplementary table 3.**

---

**Figure S1:**
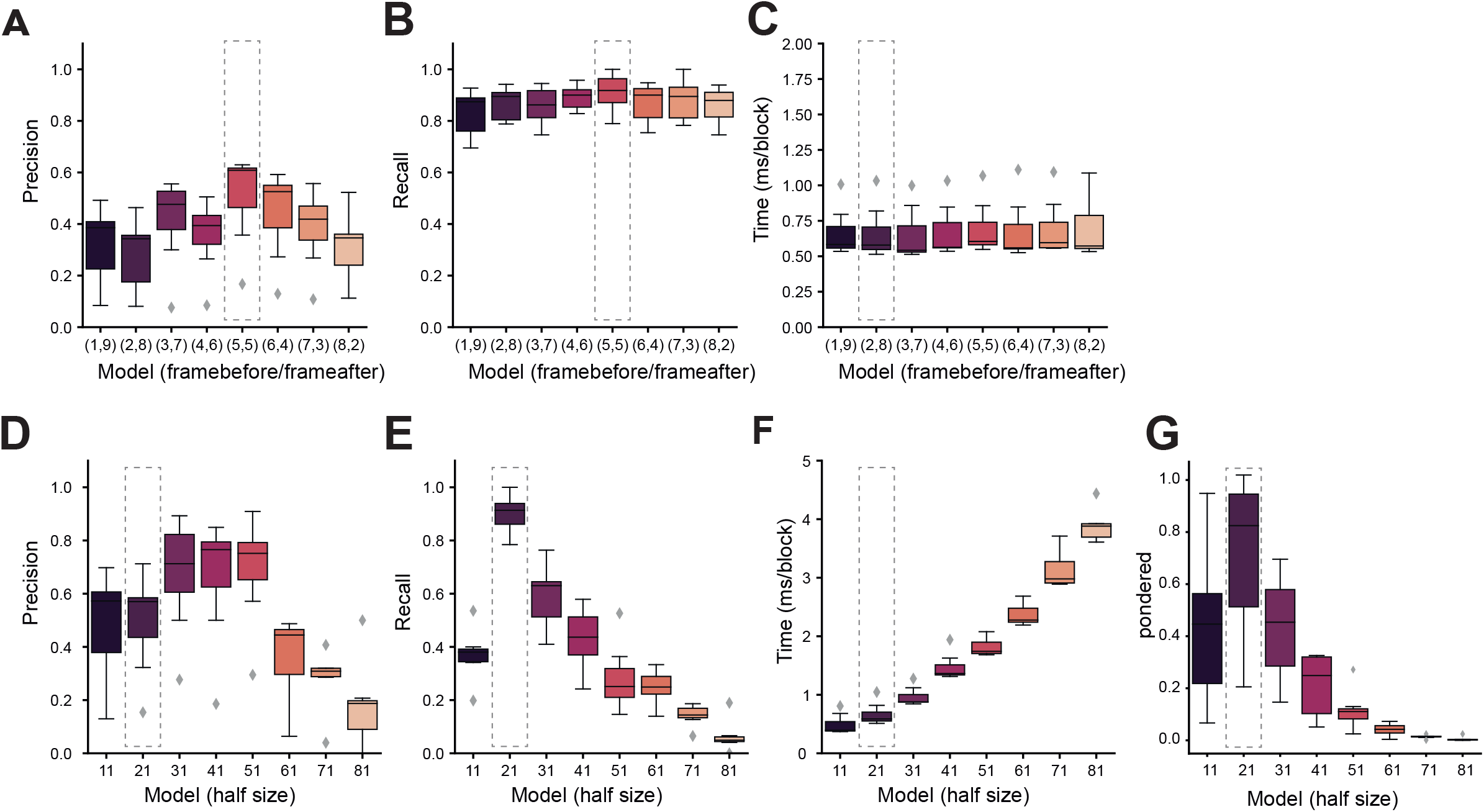
Optimisation of the search window size for extrusion detection. **A-C:** Optimisation of the temporal positioning of the search window according to the termination of extrusion (manually detected, end of apical area closure), number of frames before or after the extrusion detection point. For each position, the Precision (**A**), Recall (**B**), and time of calculation (ms per block) (**C**), were estimated on the test dataset. The optimum was obtained for a search window centered on extrusion termination (5,5). Box plots show the median, the first and third quartile. Top and bottom bars are the maximal and minimal value. Diamonds are outliers. **D-F:** Optimisation of the size (x y) of the square search window (half size in pixel after rescaling, 1 pixel=0.275 μm). For each position, the Precision (**D**), Recall (**E**), and time of calculation (ms per block) (**F**), were estimated on the training dataset. We computed then a ponderated parameter (**G**, see **Methods**) which takes into account precision, recall and calculation time which peaks for 21px. Box plots show the median, the first and third quartile. Top and bottom bars are the maximal and minimal value. Diamonds are outliers.

**Figure S2:**
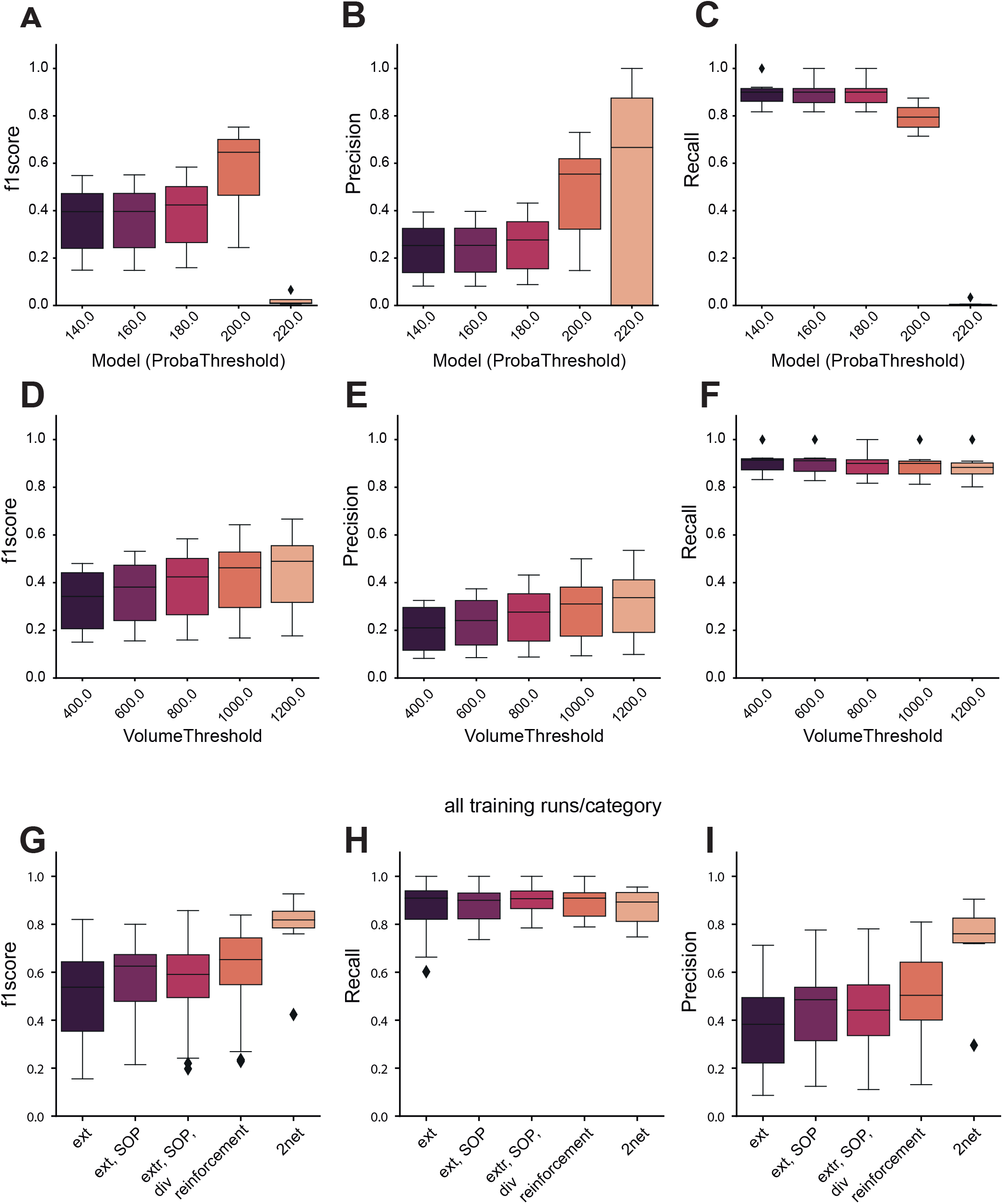
Optimisation of probability thresholding for extrusion detection and detection of new categories of cellular events. **A-C:** Optimisation of the probability threshold used to assign an extrusion event. For each threshold, the f1 score (**A**), precision (**B**), and recall (**C**), were estimated on the test dataset. The optimum f1 score was obtained for 200 (a.u.), however we used 180 in all the rest of our pipeline as we wanted to maximise the recall. Box plots show the median, the first and third quartile. Top and bottom bars are the maximal and minimal value. Diamonds are outliers. **D-F:** Optimisation of the threshold probability volume (x-y-t) used to detect extrusion event (in voxel, xy pixel=0.275μm, t=5 minutes). For each volume, the f1 score (**D**), precision (**E**), and recall (**F**), were estimated on the test dataset. The optimium for the f1score was obtained for 1200, however we used a threshold of 800 for all the rest of the pipeline to maximise the recall. Box plots show the median, the first and third quartile. Top and bottom bars are the maximal and minimal value. Diamonds are outliers. **G-I:** Changes in the model to optimise its prediction scores on extrusion. **G:** f1-score. **H:** Recall, **I:** Precision, with the initial two class model (ext), the inclusion of SOPs (ext, SOP), the inclusion of SOPs and cell divisions (ext, SOP, div), including the 3 cellular events and reinforcement (see below), and using two independent networks (2net). The results shown are the compiling of the prediction of all the testing dataset processed through 4 independent trained networks (except for 2net, which used only one pair of networks). Box plots show the median, the first and third quartile. Top and bottom bars are the maximal and minimal value. Diamonds are outliers.

**Figure S3:**
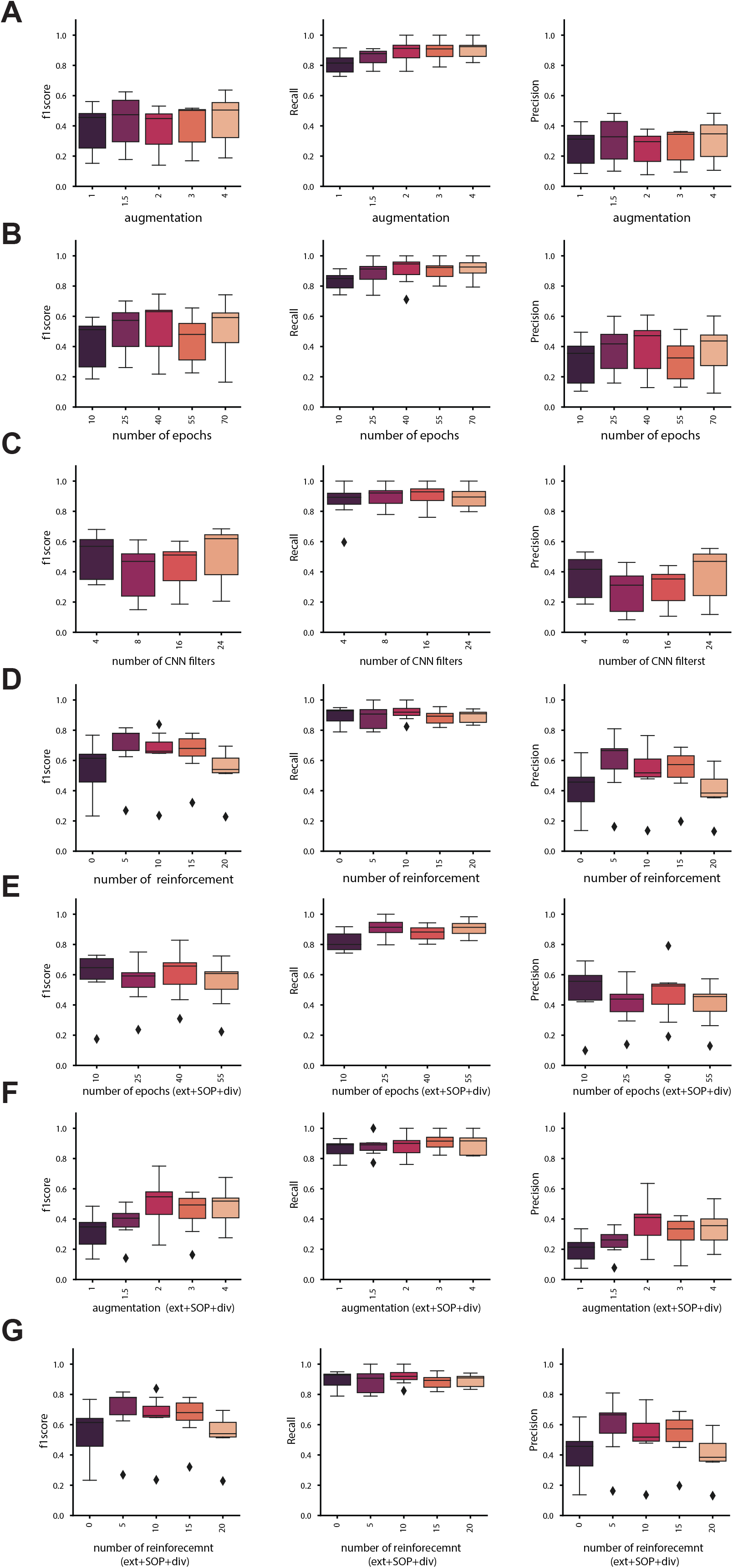
Optimisation of the hyperparameters of DeXNet. **A-F:** Optimisation of the hyperparameters of DeXtrusion to maximise the precision and the recall. For each set of parameters, the f1 score, Recall and Precision were estimated on the test dataset either for the DeXNet trained only for extrusion detection (**A-D**) or the DeXNet detecting the four categories of events (extrusion, SOPs, division, control) (**E-G**). The parameters used in this study are summarized in the **Supplementary table 3**. These parameters include the number of augmentations of the data (**A, F**), the number of epochs for model fitting (**B, E**), the number of CNN filters (**C**), and the number of reinforcements (**D, G**). Box plots show the median, the first and third quartile. Top and bottom bars are the maximal and minimal value. Diamonds are outliers.

## Movie legends

**Movie 1: Probability map of extrusions, SOPs and divisions in a WT pupal notum**

Local projection of a pupal notum expressing E-cad::GFP (grey) overlayed with the probability map of detection of extrusions (yellow), cell divisions (magenta) and SOPs (cyan). E-cad channel is shown separated on the right. Scale bar=30μm.

**Movie 2: Probability map of extrusions, SOPs and divisions in an EGFR depleted pupal notum**

Local projection of a pupal notum depleted for EGFR *(pnr-Gal4, UAS-EGFRdsRNA)*expressing E-cad::GFP (grey) overlayed with the probability map of detection of extrusions (yellow), cell divisions (magenta) and SOPs (cyan). E-cad channel is shown separated on the right. Scale bar=30μm.

**Movie 3: Probability map of extrusions, SOPs and divisions in a pupal wing**

z-projection of a WT pupal wing expressing E-cad::GFP (grey) from (Etournay et al., 2015), overlayed with the probability map of detection of extrusions (yellow) and cell divisions (magenta). E-cad channel is shown separated on the bottom. Scale bar=50μm. Note that we only took in consideration the probability overlapping the wing.

**Movie 4: Probability map of extrusions in the larval epidermal cells of the pupal abdomen**

z-projection of a WT pupal abdomen expressing E-cad::GFP (grey) from (Davis et al., 2022) overlayed with the probability map of detection of extrusions (yellow). E-cad channel is shown separated on the bottom. Scale bar=50μm. Note that we only used the prediction in the LECs and ignored the histoblasts (small cells in the clusters on the left). The cell scale used for this prediction is suboptimal for histoblasts.

## Notes

https://gitlab.pasteur.fr/gletort/dextrusion

https://doi.org/10.5281/zenodo.7586394

### Competing Interest Statement

The authors have declared no competing interest.

